# Modeling the hemodynamic response function using simultaneous EEG-fMRI data and convolutional sparse coding analysis with rank-1 constraints

**DOI:** 10.1101/2020.09.09.290296

**Authors:** Prokopis C. Prokopiou, Michalis Kassinopoulos, Alba Xifra-Porxas, Marie-Hélène Boudrias, Georgios D. Mitsis

## Abstract

Over the last few years, an increasing body of evidence points to the hemodynamic response function as an important confound of resting-state functional connectivity. Several studies in the literature proposed using blind deconvolution of resting-state fMRI data to retrieve the HRF, which can be subsequently used for hemodynamic deblurring. A basic hypothesis in these studies is that relevant information of the resting-state brain dynamics is condensed in discrete events resulting in large amplitude peaks in the BOLD signal. In this work, we showed that important information of resting-state activity, in addition to the larger amplitude peaks, is also concentrated in lower amplitude peaks. Moreover, due to the strong effect of physiological noise and head motion on the BOLD signal, which in many cases may not be completely removed after preprocessing, the neurophysiological origin of the large amplitude BOLD signal peaks is questionable. Hence, focusing on the large amplitude BOLD signal peaks may yield biased HRF estimates. To define discrete events of neuronal origins, we proposed using simultaneous EEG-fMRI along with convolutional sparse coding analysis. Our results suggested that events detected in the EEG are able to describe the slow oscillations of the BOLD signal and to obtain consistent HRF shapes across subjects under both task-based and resting-state conditions.

## 1. Introduction

Over the last 30 years, blood oxygenation level-dependent functional magnetic resonance imaging (BOLD-fMRI) has been widely used for studying brain function and its organization into functional networks (Belliveau et al., 1991; Kwong et al., 1992; Ogawa et al., 1990b, 1990a). The popularity of this technique derives from its ease to operate, non-invasive nature and high spatial resolution (Glover, 2011). The BOLD contrast mechanism depends on the dynamics of the local concentration in deoxygenated hemoglobin. The latter, in its turn, is influenced by local changes in cerebral blood flow (CBF), cerebral metabolic rate of oxygen (CMRO2), and cerebral blood volume (CBV), which are induced by increases in neuronal activity. Hence, BOLD-fMRI is an indirect measurement of neuronal activity through a series of physiological events that are collectively known as the hemodynamic response (Buxton, 2009).

This complex link between neuronal activation and its corresponding changes in BOLD-fMRI is typically modelled with the hemodynamic response function (HRF) (Buxton et al., 2004). A large number of studies in the literature pointed out that the HRF is region- and subject-specific (Aguirre et al., 1998; Handwerker et al., 2004; Rangaprakash et al., 2017), and that more accurate shapes of HRF are needed to obtain precise localization of brain activity (Lindquist et al., 2009b; Lindquist and Wager, 2007; Loh et al., 2008). In addition, there is a growing body of evidence to suggest that functional or effective connectivity measures suffer from the sluggishness of the HRF, and trying to account for the hemodynamic blurring using inaccurate HRF shapes make unclear whether observed changes in connectivity are due to neuronal activity or HRF variability (Deshpande et al., 2010; Handwerker et al., 2012; Rangaprakash et al., 2018; G.-R. Wu et al., 2013).

HRF estimation has been a topic of continual research since the inception of BOLD-fMRI. The first class of HRF estimation algorithms that are found in the literature includes parametric identification methods, which assume a specific structure for the unknown HRF. In this context, the HRF shape is controlled by a few parameters that could be estimated from the data. Algorithms of this type usually involve Gaussian HRF shapes (Kruggel and Cramon, 1999; Rajapakse et al., 1998), gamma HRF shapes (K. J. J. Friston et al., 1998; Miezin et al., 2000), or spline-like functions (Gössl et al., 2001). The second class includes non-parametric methods, which make no prior hypotheses about the shape of the HRF estimates. Such methods include selective averaging (Dale and Buckner, 1997), smooth HRF filtering (Goutte et al., 2000), Bayesian methods (Ciuciu et al., 2003; Marrelec et al., 2003b), linear subspace methods (Hossein-Zadeh et al., 2003; Steffener et al., 2010; Woolrich et al., 2004b), wavelet methods (Lina et al., 2010), and machine learning methods (Güçlü and van Gerven, 2017; Luo and Puthusserypady, 2007; Pedregosa et al., 2015). Most of these HRF estimation methodologies were developed under the assumption that neuronal activity and the corresponding BOLD responses are known. In task-related studies, neuronal activity is assumed to closely follow the external stimulus or task execution. In resting-state studies on the other hand, the absence of a specific task renders HRF estimation a challenging endeavor.

Recent studies in the literature attempted to address HRF estimation in resting-state fMRI using events detected in the BOLD signal (G.-R. Wu et al., 2013). These works laid on the hypothesis that the neural events that govern the dynamics of the brain in resting-state are reflected in the large amplitude peaks or transients in the BOLD signal, which can be retrieved using point-process analysis (PPA) (Tagliazucchi et al., 2012, 2011) or sparse-promoting deconvolution (Caballero Gaudes et al., 2013; Karahanoğlu et al., 2013). Subsequently, resting-state fMRI was considered as spontaneous event-related, and the HRF was estimated using those pseudo-events (Abe et al., 2015; Alavash et al., 2016; Case et al., 2017; Chen and Glover, 2015; Iwabuchi et al., 2014; Rangaprakash et al., 2018; G. Wu et al., 2013). Spontaneous, large amplitude peaks observed in the BOLD have also been associated with functional resting-state networks (Karahanoğlu et al., 2013; Petridou et al., 2013; Tagliazucchi et al., 2012), as well as transient, recurrent patterns of co-activation observed with fMRI (Liu et al., 2013; Liu and Duyn, 2013), suggesting their neurophysiological origins.

Large amplitude peaks or transients in the BOLD, however, in addition to neural events may also reflect motion (Power et al., 2012) and physiological noise, such as spontaneous fluctuations in arterial CO2 (Golestani et al., 2016; Prokopiou et al., 2019, 2016), cardiac pulsatility (Glover et al., 2000), respiration and heart rate variability (Birn et al., 2008; Chang et al., 2013, 2009; Kassinopoulos and Mitsis, 2019). These non-neuronal sources of BOLD signal variability have been shown to elicit networks of coherent BOLD activity, which resemble previously reported resting-state networks derived from fMRI data (Chen et al., 2019; Nalci et al., 2019; Nikolaou et al., 2016; Shokri-Kojori et al., 2018). In addition, covariation of the BOLD signal in different brain regions that is sufficient to give rise to spatial patterns of resting-state activity can be also observed at the timings of lower amplitude peaks, or even at regularly or randomly selected timepoints along the time course of the signal. Hence, the neural events that govern the dynamics of the brain in resting-state may not be reflected only in the large amplitude peaks of the BOLD signal. In light of the above considerations, the neurophysiological origin of its high amplitude peaks is questionable. A propitious avenue for obtaining more reliable HRF estimates from resting-state measurements is by using multimodal imaging techniques, such as simultaneous electro-encephalography (EEG)-fMRI, where the neuronally-driven activity in the EEG is combined with the high spatial resolution of the fMRI.

In this work, we initially employed resting-state fMRI data and PPA analysis to define sparse events in the BOLD signal corresponding to large amplitude peaks. We showed that the mean interval between the detected events is smaller than the maximum (Nyquist) sampling interval that is required to retain the slow dynamics of the BOLD during the resting state. Subsequently, using seed-based correlation analysis with a seed selected in the precuneus cortex (PCC) we showed that the covariation between regularly spaced, as well as randomly spaced BOLD samples, which do not always coincide with high amplitude peaks obtained from individual voxels and the seed is sufficient to derive the default mode network (DMN) of the brain. Moreover, after regressing out the defined PPA events from the original BOLD time-series, we showed that new events obtained by re-applying PPA on the residual time-series yield the same patterns of concurrent activity between individual voxels and the seed (conditional rate maps) as the ones obtained using the events defined in the original time-series. Therefore, we concluded that the important information of resting-state BOLD activity is not condensed only in its high amplitude peaks, and that the neurophysiological origin of high amplitude peaks in the BOLD is not warranted. Hence, using these events for HRF estimation may yield biased estimates.

To define more reliable neural events that can be used for HRF estimation, we employed EEG data collected simultaneously with BOLD-fMRI along with convolutional sparse coding (CSC) analysis with rank-1 constraints. CSC analysis is a dictionary learning technique that can be used to provide information about the spatial pattern, temporal waveform, and the timing of neural events defined in EEG data. We initially performed this analysis using task-based data collected during two separate experiments: (1) a visual target detection and (2) a hand grip task. To show the functional relevance of the detected CSC events for each task we performed event-related fMRI analysis. Subsequently, we compared the resultant activation maps with the corresponding maps obtained using external measurements of the subjects’ behavioral response to each task. Our results revealed concordance between the activation maps obtained in each case, suggesting that CSC analysis can be used to detect events in the EEG that are associated with each task, and which are able to describe the slow dynamics of the BOLD signal. In addition, we used the detected CSC events to obtain estimates of the HRF in large functionally defined ROIs, which revealed consistent shapes across subjects.

We also employed CSC analysis to define events in resting-state EEG data collected simultaneously with BOLD-fMRI. The results suggested that CSC analysis can be used to detect events in the EEG even during resting-state, where SNR is lower, and that these events can be used to obtain reliable estimates of the resting-state HRF. The latter could be of great importance for hemodynamic deblurring in resting-state fMRI connectivity studies.

## 2. Methods

### 2.1. Experimental methods

Two datasets were employed in this study. Dataset 1: Seventeen healthy subjects (6 females; mean of 27.7 years, 20-40 range) participated in a visual target detection task (visual oddball paradigm). All subjects gave informed consent following the protocol of the Columbia University Institutional Review Board^1^. Dataset 2: 12 healthy volunteers (age range 20-29 years) participated in a resting-state experiment (session 1) followed by a hand grip task (session 2) after giving a written informed consent in accordance with the McGill University Ethical Advisory Committee. All participants were right-handed according to the Edinburgh Handedness Inventory (Oldfield, 1971).

### 2.2. Experimental paradigm

#### *Dataset 1* (Figure 1a)

A total of 125 stimuli were presented for 200 ms each. The inter-trial interval was uniformly distributed between 2–3 s. The standard stimuli were large red circles. The target stimuli were small green circles presented with probability 0.2. Both visual cues were presented to subjects on isoluminant gray backgrounds (3.45° and 1.15° visual angles) using the E-Prime software (Psychology Software Tools) and VisuaStim Digital System (Resonance Technology). The first two stimuli were constrained to be standards. The subjects were asked to respond to target stimuli, using a button press with the right index finger on an MR-compatible button response pad. Response time (RT) events were modeled using unit amplitude boxcars with onset at stimulus time and offset at subject’s response time indicated by the button press.

**Figure 1.**
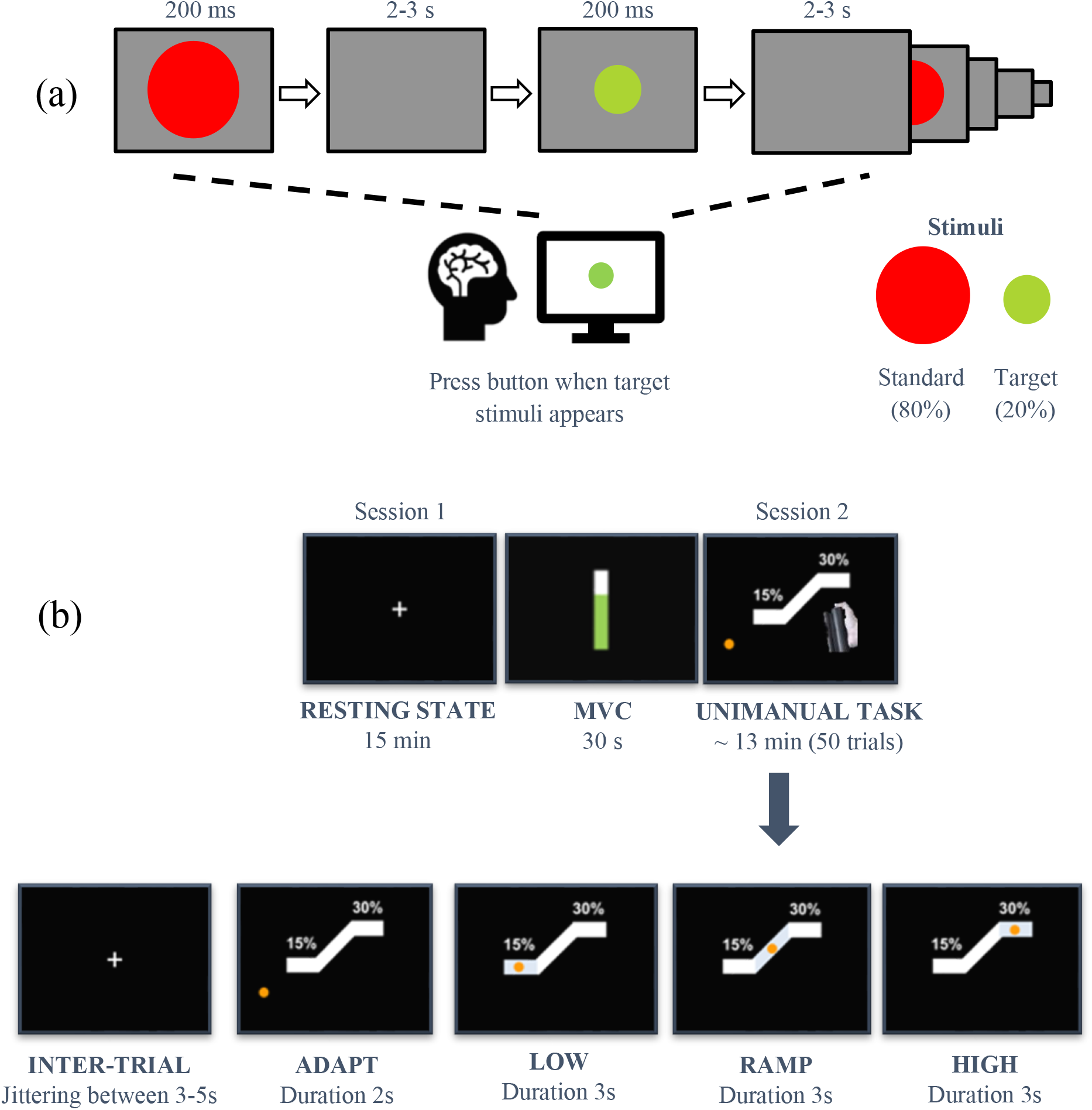
(a) Schematic representation of the visual oddball paradigm that was employed for the collection of dataset 1. (b) Illustration of the paradigm that was employed for the collection of dataset 2.

#### *Dataset 2* (Figure 1b)

The study was divided in two scans. In the first scan, subjects were instructed to stare at a white fixation cross hair displayed in a dark background (resting-state experiment). After the first scan, the maximum voluntary contraction (MVC) was obtained from each subject using the same hand gripper that was employed during the hand grip task. In the second scan, subjects were asked to perform unimanual isometric right-hand grips to track a target as accurately as possible while receiving a visual feedback. The task consisted of 50 trials, and each trial lasted 11 s. At the beginning of each trial, an orange circle appeared on the screen and subjects had to adapt their grip force at 15% of their MVC to reach a white vertical block (low force level). This force was held for 3 s. Subsequently, subjects had to linearly increase their force following a target block to reach 30% of their MVC over a 3-s period, and to hold their grip force at this level for another 3 s (high force level). The inter-trial interval was randomly jittered between 3-5 s.

### 2.3. EEG data acquisition and preprocessing

#### Dataset 1

EEG was continuously recorded during the fMRI scanning at 1 kHz using a custom-built, multi-channel MR-compatible EEG system (Goldman et al., 2009), with differential amplifier and bipolar EEG cap. The caps were configured with 36 Ag/AgCl electrodes (including the left and right mastoids), arranged as 43 bipolar pairs. The gradient artifact was removed via mean template subtraction. Subsequently, a 10 ms median filter was applied to remove residual spike artifacts. The gradient-free data were then re-referenced to a 34-electrode space (Walz et al., 2013). Further preprocessing was performed using 1 Hz high pass filtering to remove DC drift, notch filtering at 60 and 120 Hz to remove powerline artifacts, 70 Hz low pass to remove high frequency artifacts and down sampling at 150 Hz. Temporal independent component analysis (ICA) (Delorme and Makeig, 2004) was performed on each subject separately and non-neural sources of noise were removed using MARA (I. Winkler et al., 2014). After preprocessing, one subject was excluded from further analysis due to excessive noise that remained in the data.

#### Dataset 2

EEG was continuously recorded during the fMRI scanning at 5 kHz using a 64 channel MR-compatible EEG system. The caps were configured with ring Ag/AgCl electrodes, which were distributed according to the 10/20 system and referenced to electrode FCz (Brain Products GmbH, Germany). EEG data acquired inside the scanner were corrected off-line for gradient and ballisto-cardiogram (BCG) artifacts using the BrainVision Analyser 2 software package (Brain Products GmbH, Germany). The gradient artifact was removed via adaptive template subtraction (Allen et al., 2000). Gradient-free data were band-passed from 1-200 Hz, notch-filtered at 60, 120, and 180 Hz to remove power-line artifacts, and down-sampled to a 400 Hz sampling rate. The BCG artifact was removed as follows: First, temporal independent component analysis (ICA) was performed on each subject separately (Delorme and Makeig, 2004). Temporal ICA components associated with spatial patterns corresponding to BCG artifacts were visually inspected to identify the one exhibiting periodic peaks at around 1 Hz frequency, and thus be more likely to be associated to heartbeats, while accounting for most of the variance in the data. Subsequently, this component was used as a surrogate of the ECG signal to detect heartbeat events. Lastly, the BCG artifact was removed via adaptive template subtraction on a channel-by-channel basis using the detected heartbeat events (Allen et al., 1998). Subsequently, the data were re-referenced to average reference, and a second temporal ICA was performed. Noisy components associated with non-neural sources were detected and removed using MARA (I. Winkler et al., 2014). Lastly, the noise free data were down sampled to a 100 Hz sampling rate. After preprocessing, one subject was excluded from further analysis due to excessive noise that remained in the data.

### 2.4. Hand grip force measurements

For dataset 2, a non-magnetic hand clench dynamometer (Biopac Systems Inc, USA) was used to measure the subjects’ hand grip force strength during the execution of the hand grip task (session 2). The dynamometer was connected to an MR compatible Biopac MP150 data acquisition system from which the signal was transferred to a computer.

### 2.5. BOLD acquisition and preprocessing

#### Dataset 1

One hundred seventy echo-planar imaging (EPI) functional volumes were acquired on a 3T Philips Achieva MR Scanner (Philips Medical systems). EPI sequence parameters: TR/ TE =2000/35 ms (Repetition/Echo Time), Voxel size = 3×3×4 mm^3^, 32 slices with 64×64 voxels, 3 mm in-plane resolution, Slice thickness = 4 mm, and 0 mm gap. For each subject, a single 1×1×1 mm^3^ spoiled gradient recalled (SPGR) image was also acquired for purposes of registration.

#### Dataset 2

Whole-brain BOLD-fMRI volumes were acquired on a 3T MRI scanner (Siemens MAGNETOM Prisma fit) with a standard T2*-weighted echo planar imaging (EPI) sequence. Sequence parameters: TR/TE = 2120/30 ms (Repetition/Echo Time), Voxel size = 3×3×4 mm^3^, 35 slices, Slice thickness = 4 mm, Field of view (FOV) = 192 mm, Flip angle = 90°, Acquisition matrix = 64×64 (RO×PE), Bandwidth= 2368 Hz/Px. For each subject, a single 1×1×1 mm^3^ magnetization-prepared rapid gradient-echo (MPRAGE) high-resolution T1-weighted structural image was also acquired for the purposes of registration.

For both datasets, fMRI data preprocessing was carried out using the FSL Software Library (FMRIB, Oxford, UK, version 5.0.10) (Jenkinson et al., 2012). fMRI preprocessing steps included image realignment, spatial smoothing with a Gaussian kernel of 5 mm full-width at half maximum (FWHM), and high-pass temporal filtering. Spatial ICA was carried out for each subject using MELODIC, and spatial maps associated with cardiac pulsatility, head motion, susceptibility and other MRI-related artefacts were removed. Noise-free data were subsequently registered with T1-weighted structural images and normalized to the Montreal Neurological Institute (MNI)-152 brain template, with resolution of 2×2×2 mm^3^.

### 2.6. Data analysis

#### 2.6.1. High amplitude peaks in resting-state BOLD correspond to sub-Nyquist sampling intervals

In this analysis we sought to investigate the following hypotheses: (1) the interval between events defined at the high amplitude peaks in resting-state BOLD is smaller than half the Nyquist interval of the signal, (2) the covariation of regularly- or randomly-spaced BOLD samples obtained with a sampling interval smaller than the Nyquist interval of the BOLD is sufficient to derive patterns of resting-state brain activity, and (3) the patterns of coactivation (conditional rate maps) obtained using events defined at lower amplitude BOLD signal peaks are similar to the ones obtained with events defined at higher amplitude peaks. These hypotheses suggest that relevant information related to the spatiotemporal dynamics of resting-state activity, in addition to the high amplitude peaks in the BOLD may also be condensed in lower amplitude peaks. Moreover, taking into consideration the strong effects of physiological processes in the BOLD signal, they also suggest that the neurophysiological origin of the high amplitude peaks in the BOLD is not warranted.

##### Group average power spectral density (PSD)

The Nyquist interval of the resting-state BOLD-fMRI signal was determined based upon the group average PSD obtained using the resting-state data from dataset 2.

A PSD was estimated at each voxel using the Welch method: the original data were segmented into 8 data segments on which a Hamming window was applied. Consecutive data segments were overlapped by 50%. For each of the 8 segments the Fourier Transform (FFT) was calculated and the power of the FFT coefficients was averaged across the overlapping windows. The PSD obtained at each voxel was initially averaged within subjects, and subsequently between subjects to obtain a group average PSD.

##### Point process analysis (PPA)

Events corresponding to large amplitude peaks in the BOLD were defined using PPA (Tagliazucchi et al., 2012). The BOLD time-series at each voxel was initially normalized by its own standard deviation (SD). Subsequently, an event was defined every time the signal crossed a threshold (1 SD) from below.

##### Seed-based Pearson correlation analysis

Seed-based correlation analysis was performed between individual voxels and a seed selected in the precuneus cortex (PCC). The seed time course consisted of the averaged signal from all voxels within a ball of radius 10 mm, centered at the MNI voxel co-ordinates X = 48, Y = 37, Z = 56. The data were down-sampled using down-sampling factors of 8, 10, 12, and 14 points. These values were defined based on the 25th and 75th percentile of the boxplot of voxel-wise mean PPA sampling interval values obtained for each subject, which are shown in Figure 3b. A random down-sampling was also performed, where the sampling interval between any two consecutive samples was determined based on a uniform distribution. The minimum and maximum values of the distribution were defined based on the 25th and 75th percentile of the boxplots shown in Figure 3b. BOLD signal correlations between down-sampled time-series from individual voxels and the seed were obtained using the Pearson correlation coefficient, which is given by

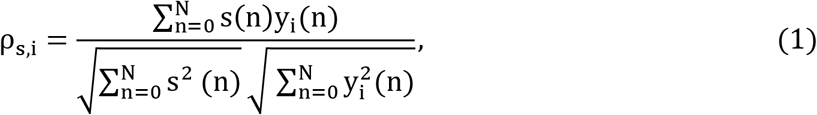

where s(n) denotes the time-series of the seed, and y_i_(n) the BOLD time-series of voxel i, at time n. Correlation scores were converted to z-scores using the Fisher z-transform, given by

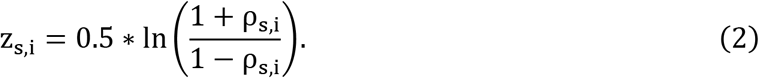

Individual correlation maps were warped to the 2 mm^3^ MNI template and fed into the second-level analyses with a voxel-wise one-sample t-test to compare functional connectivity between individual voxels and the seed. Group-level t-maps are shown in Figure 4. In each case, p-values were converted into a False Discovery Rate (FDR), and the statistical maps were thresholded at p_FDR_ < 0.005.

##### Residual analysis of PPA events

To investigate the co-activation patterns of lower amplitude peaks in the resting-state BOLD we proceeded as follows:

First, large amplitude BOLD signal peaks were detected using PPA, and subsequently regressed out from the original BOLD time-series to obtain the residual time-series. Specifically, a BOLD prediction 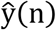 was obtained at each voxel as

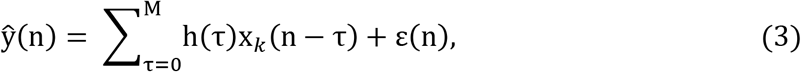

where x_k_(n − τ) is a k-lagged version of the PPA events defined on the original BOLD time-series, and h(τ) an estimate of the resting-state HRF, which was obtained using a blind deconvolution method proposed by (G.-R. Wu et al., 2013). Then, the residual time-series at each voxel were calculated as

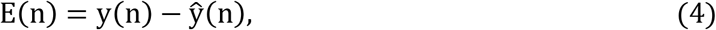

where y(n) denotes the original BOLD time-series.

Subsequently, PPA analysis was applied on the residual time-series E(n), and a conditional rate map between individual voxels and a seed (PCC) was constructed (see Conditional rate maps below).

This procedure was repeated four times: the first time, PPA events were defined based on the original resting-state BOLD-fMRI data. The other three times, PPA events were defined based on the residual data obtained in the previous iteration. The conditional rate maps obtained in each case are shown in Figure 5.

##### Conditional rate maps (Tagliazucchi et al., 2012)

PPA events were defined for both the seed and individual voxels in the brain (targets). Every time a PPA event at a target voxel was defined up to 2-time steps later than in the seed, the rate at the target was increased by one unit. Lastly, this rate was normalized by the number of points in the seed.

#### 2.6.2. Convolutional sparse coding (CSC) analysis

To define transient events in the EEG data we employed a multivariate CSC with rank-1 constraint (Jas et al., 2017; La Tour et al., 2018), which is described by

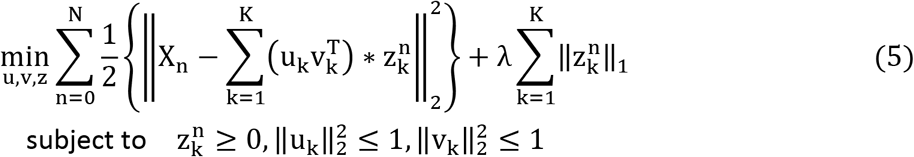

where X_n_ ∈ ℝ^P×N^ denotes the EEG timeseries measured from P sensors at time n = 0, …, N, λ > 0 the regularization parameter, u_k_ ∈ ℝ^P^ the k-th spatial pattern 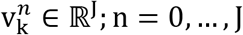 the k-th temporal waveform, and 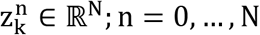 the sparse vector of EEG events associated with the k-th temporal waveform. The rank-1 constraint is consistent with Maxwell’s equations and the electromagnetic properties of the brain waves, which propagate instantaneously inside the head volume and add up linearly at the level of EEG sensors (Hari and Puce, 2017).

We hypothesized that the duration of the temporal waveforms is 1 s. Therefore, J was set equal to 150, and 100 for dataset 1, and 2, respectively. The regularization parameter λ > 0 controls the sparsity of 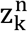 induced by the *l*_1_-norm: the higher the regularization parameter λ, the higher the sparsity. In this work, λ was set equal to 0.08 for dataset 1, and 0.1 for dataset 2. These values were determined empirically for each dataset, as they were found to balance between sparsity and BOLD prediction accuracy. The optimization problem described by equation (5) was solved efficiently using a locally greedy coordinate descent algorithm (Moreau et al., 2018), as well as precomputation steps for faster gradient computations as described in (La Tour et al., 2018).

The number of rank-1 atoms that can be obtained from a given EEG dataset using CSC analysis is equal to the total number of the EEG sensors. However, the rank of the data may have been decreased during preprocessing due to application of ICA and removal of noise-related ICA components. To account for this reduction in the dimensionality of the data the maximum number of CSC atoms that was estimated for each subject was equal to the number of ICA components that were retained in the data (Prokopiou and Mitsis, 2019). Moreover, some of the CSC atoms exhibited spatial patterns of typical EEG artifacts observed during EEG-fMRI experiments, such as gradient, BCG, and eye-blink artefacts (Figure 6). These patterns were isolated and subsequently removed from any subsequent analysis.

To obtain a total event time-series from the 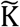 selected CSC spatiotemporal atoms (Figure 2), each special pattern u_k_ ∈ ℝ^P^was initially multiplied with its associated event time-series 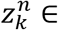 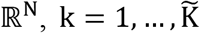. This yielded a set of 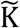 rank-1 matrices of event time-series 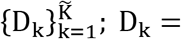 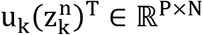, 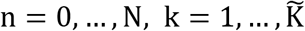, which were projected to the EEG sensor level by taking the sum over all selected atoms

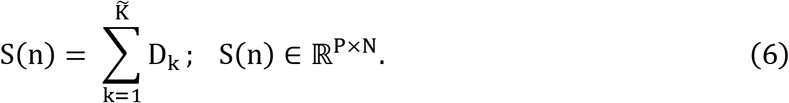

**Figure 2.**
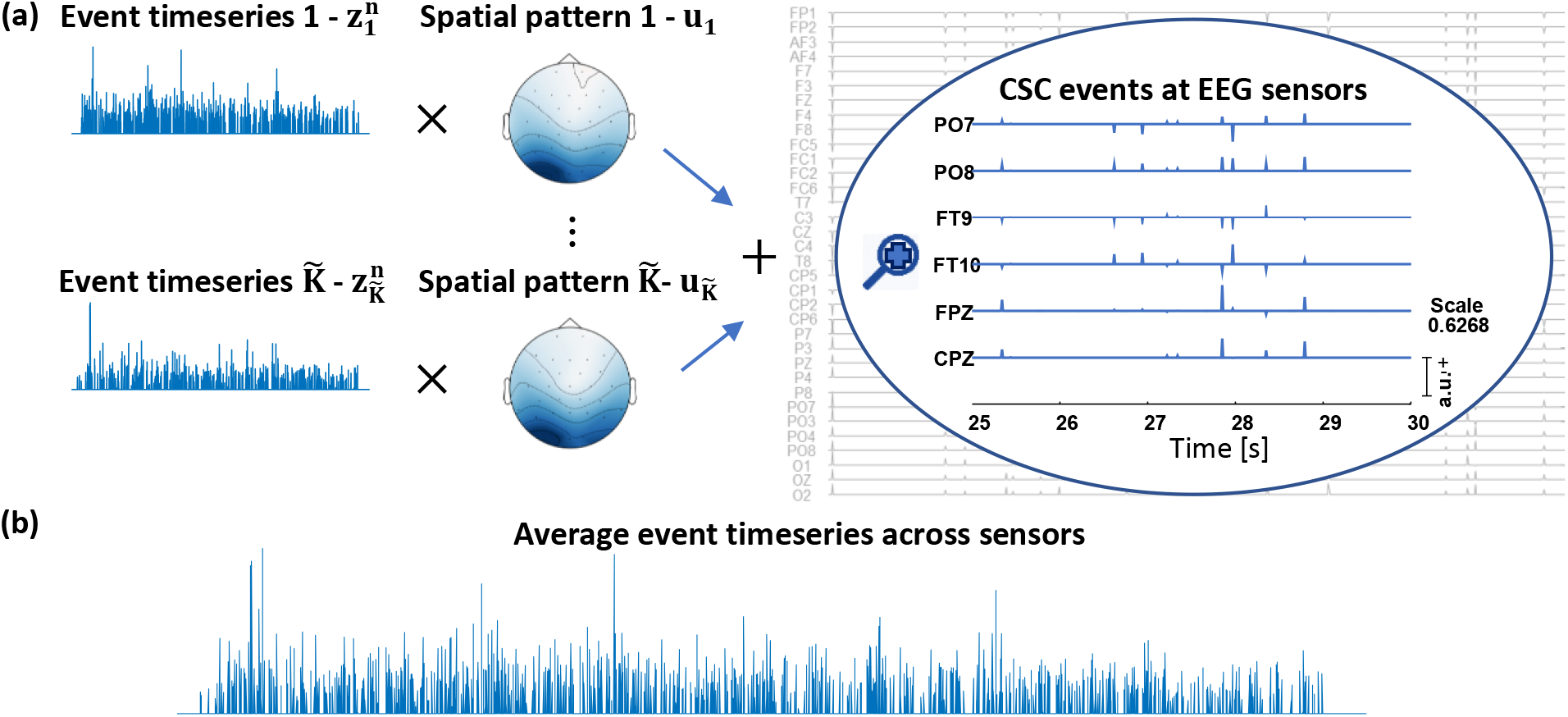
Construction of a total CSC event time-series from the individual CSC atoms. **(a)** The event time-series 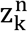 of the k-th CSC atom is multiplied with its associated spatial pattern uk, 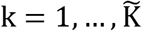. This results into a rank-1 matrix 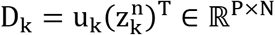 of event time-series, which is associated with the k-th CSC atom. The individual event time-series are projected at the EEG sensor level by taking the sum of all the 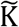 rank-1 matrices D_k_. **(b)** A total event time-series is obtained as the mean of the projected CSC events across all EEG sensors.

A unique event time-series 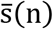 was obtained for each subject as the mean of the reconstructed events at the sensor level, given by

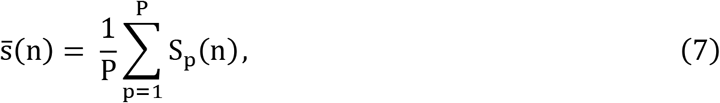

where P denotes the total number of EEG sensors.

#### 2.6.3. Voxel-wise analysis

##### Dataset 1

We initially employed dataset 1 along with CSC analysis to define events associated with the visual target detection task. Subsequently, we used the mean CSC event time-series 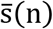 to obtain BOLD predictions in a voxel-wise fashion using a standard event-related fMRI analysis. BOLD predictions were also obtained using the subjects’ behavioral response time (RT) events.

CSC and RT events were convolved with the canonical, double gamma HRF, which is implemented in SPM (https://www.fil.ion.ucl.ac.uk/spm/) to generate one regressor for each event type. Subsequently, a mixed-effects approach was used to model activation across subjects using FEAT (Woolrich et al., 2004a, 2001). The search for activation was contained within gray matter. This analysis was performed for each regressor independently. The resultant activation maps obtained in each case were qualitatively compared between them in order to assess the extend and region specificity of the activation patterns induced by each event type (Figure 7a).

##### Dataset 2

We employed the EEG-fMRI data collected during the hand-grip task (Figure 1b) along with CSC analysis to define sparse events in the EEG. Our aim was to investigate the dynamic interactions between these events and the BOLD signal at individual voxels using FIR model analysis.

The FIR model describes the output of a linear and time-invariant system as a weighted sum of past input values, where the weighting coefficients are given by the impulse response function. The FIR model is given by

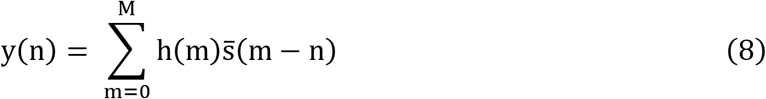

where y(n) denotes the output (i.e. the BOLD signal), and 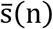 the input (i.e. the mean event time-series of the reconstructed CSC events at the sensor level) of the system. h(n) denotes the unknown impulse response (i.e. the HRF), and M the system memory.

For the estimation of the unknown HRF, we employed a function expansion technique along with a set of orthonormal basis functions (Marmarelis, 1993)

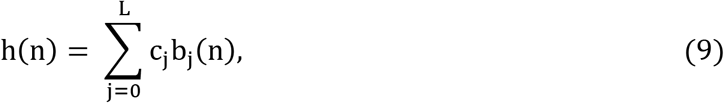

where {b_j_(n); j = 0, …, L − 1} is a set of L basis functions, and c_j_ the unknown expansion coefficients. Substitution of (9) in (8) yields

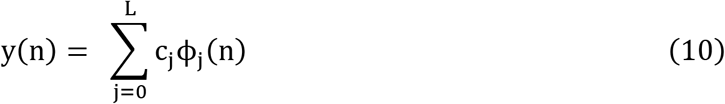

where 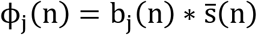. Equation (10) can be rewritten in a compact matrix form as

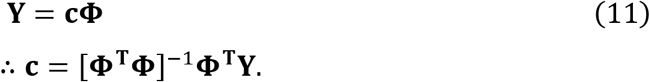

The unknown expansion coefficients **c** can be obtained using ordinary least squares regression.

An important step in the application of the function expansion technique is the selection of a proper basis set, as it influences the obtained estimates. The choice of a basis set depends on the dynamic behavior of the system to be modelled. One basis set that has been extensively used in the literature, particularly in the case of physiological systems is the Laguerre basis. The Laguerre functions are orthonormal and exponentially decaying curves, which constitute a basis for the space of square integrable functions L^2^[0, ∞]. These properties of the Laguerre functions makes them suitable for modeling causal systems with finite memory (Marmarelis, 1993).

In the present work we employed the spherical Laguerre basis set (Leistedt and McEwen, 2012). The spherical Laguerre basis is a smoother variant of the Laguerre basis, which allowed us to obtain robust HRF estimates even during resting conditions, where the signal-to-noise ratio (SNR) is low. The j-th spherical Laguerre function b_j_(n); j = 0, …, L − 1; n = 0, …, M is given by

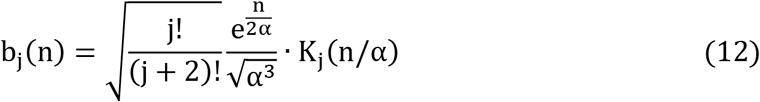

where α ∈ ℝ_+_ is a parameter that determines the rate of exponential decay of b_j_(n), and K_j_(n) is the j-th generalized Laguerre polynomial of order two, defined as

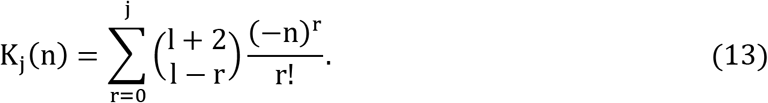

To prevent overfitting, the range for the total number L of basis functions and the range for the parameter α was selected to be 2 < L ≤ 4 and 0.5 < α < 1. Model performance was evaluated in terms of the mean-squared prediction error (MSE), which is given by

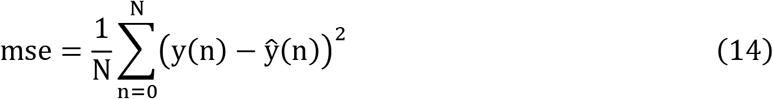

where y(n) denotes the measured BOLD, and 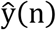 the predicted BOLD time-series obtained using equation (8). The optimal value for the structural Laguerre parameters L and α was determined in terms of the minimum MSE using a grid search.

Performing grid search to determine optimal values for the Laguerre parameters at each voxel incurs a heavy computational burden. To reduce the computational complexity for the estimation of voxel-specific HRFs, we initially obtained HRF estimates in large structurally defined ROIs for each subject, according to the Harvard-Oxford cortical atlas^2^. Subsequently, we applied singular value decomposition (SVD) on the set of ROI-specific HRFs corresponding to each subject in order to obtain a reduced set of orthonormal functions that account for the major fraction of the variability in this set. This procedure yielded a set of two singular vectors representative of the ROI-specific HRF shapes obtained from each subject, as it was found that the two absolutely largest singular values accounted for more than 80% of variability of the original set of HRFs. For all subsequent analyses, the HRF curve estimates were obtained using equations (8)–(11) along with the set of two orthonormal functions obtained for each subject.

To investigate the functional relevance of the detected CSC events, the mean event time-series 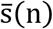 was convolved with each basis function, and a BOLD prediction was obtained at each voxel. Then, the F-score was calculated

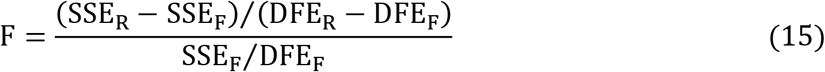

where SSE_F_ and SSE_R_ are respectively the residual sum of squares of the full and null model. DFE_F_ and DFE_R_ denote the number of degrees of freedom for the full and null model, respectively. The statistic F follows a 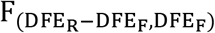 distribution. A large value of F indicates that the detected CSC events significantly contribute to BOLD signal variance. Finally, random field theory (Worsley et al., 1996) was employed to compute the significance level corrected for multiple comparisons, where search for activation was contained within gray matter.

## 3. Results

### 3.1. High amplitude peaks in the BOLD correspond to sub-Nyquist sampling intervals

The upper panel in Figure 3 shows the average PSD obtained across all subjects from the resting-state fMRI data of dataset 2. The PSD has a peak around 0.013 Hz. According to the Nyquist sampling theorem, sampling with a sampling frequency higher than 0.026 Hz, or equivalently with a sampling interval smaller than approximately 38 s is sufficient to describe the slow dynamics of the BOLD observed during the resting-state. The bottom panel in Figure 3 shows boxplots of the mean interval between high amplitude peaks in the BOLD signal at each voxel obtained using PPA, for each subject. The voxel-wise mean sampling intervals were found to be smaller than the Nyquist sampling interval, which is shown with a dashed line, for most of the subjects. This suggests that using samples obtained from the BOLD signal with any regular or irregular sampling interval smaller than the Nyquist interval, which includes the interval between its high amplitude peaks, is sufficient to preserve the signal’s slow dynamics observed during the resting-state. Since the slow dynamics of the BOLD can be described by samples obtained using any sub-Nyquist sampling interval that may not necessarily coincide with its large amplitude peaks, this suggests that the relevant information related to the dynamics of the signal observed during resting-state is not condensed only at the timings of its high amplitude peaks.

**Figure 3.**
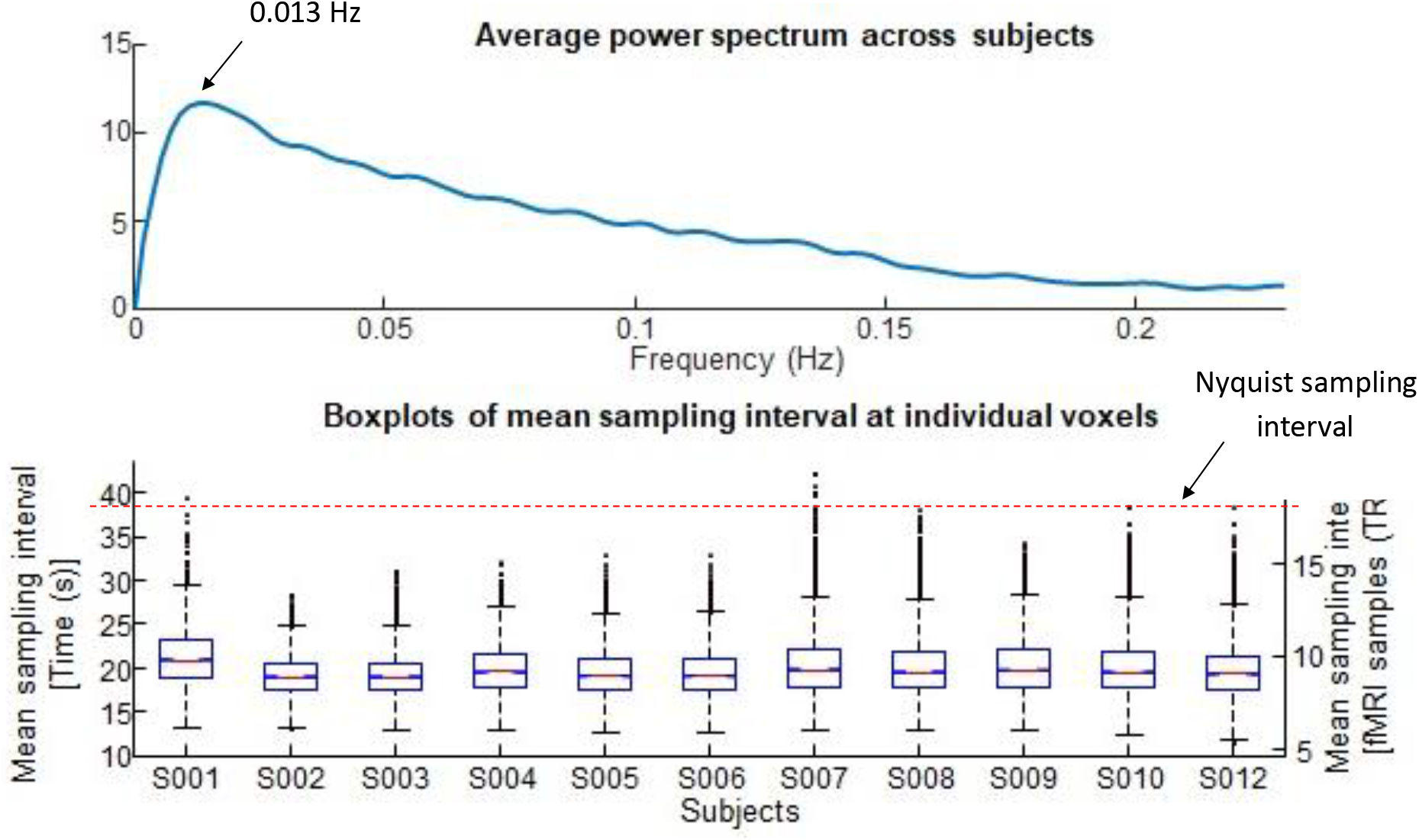
Top panel: average power spectral density (PSD) across subjects obtained from resting-state BOLD-fMRI data. The curve shows a peak around 0.013 Hz. Sampling the BOLD signal with a sampling interval smaller than approximately 38 s, or equivalently with a sampling rate higher than 0.026 Hz, preserves the slow dynamics of the BOLD signal during resting-state (Nyquist sampling theorem). Bottom panel: boxplots of mean irregular sampling intervals obtained at each voxel, for all subjects during resting-state. For most of the subjects, the voxel-wise mean sampling interval is smaller than the Nyquist sampling interval limit of 38 s (shown with a dashed line).

Figure 4a shows group level results of seed-based correlations between individual voxels and a seed selected in the precuneus cortex. The seed time-series was constructed for each subject using the mean of all voxels within a ball of radius 10 mm centered at the MNI voxel coordinates X = 48, Y = 37, Z = 56. Correlations were calculated using down-sampled time-series, where down-sampling was performed with a dawn-sampling factor of 8, 10, 12, and 14 points. These values were determined based on the 25th and 75th percentile of the voxel-wise mean sampling intervals shown in the boxplots in Figure 3. In each case, the seed-based correlations revealed connectivity in the DMN, although the extend of the network is decreased at larger sampling intervals

**Figure 4.**
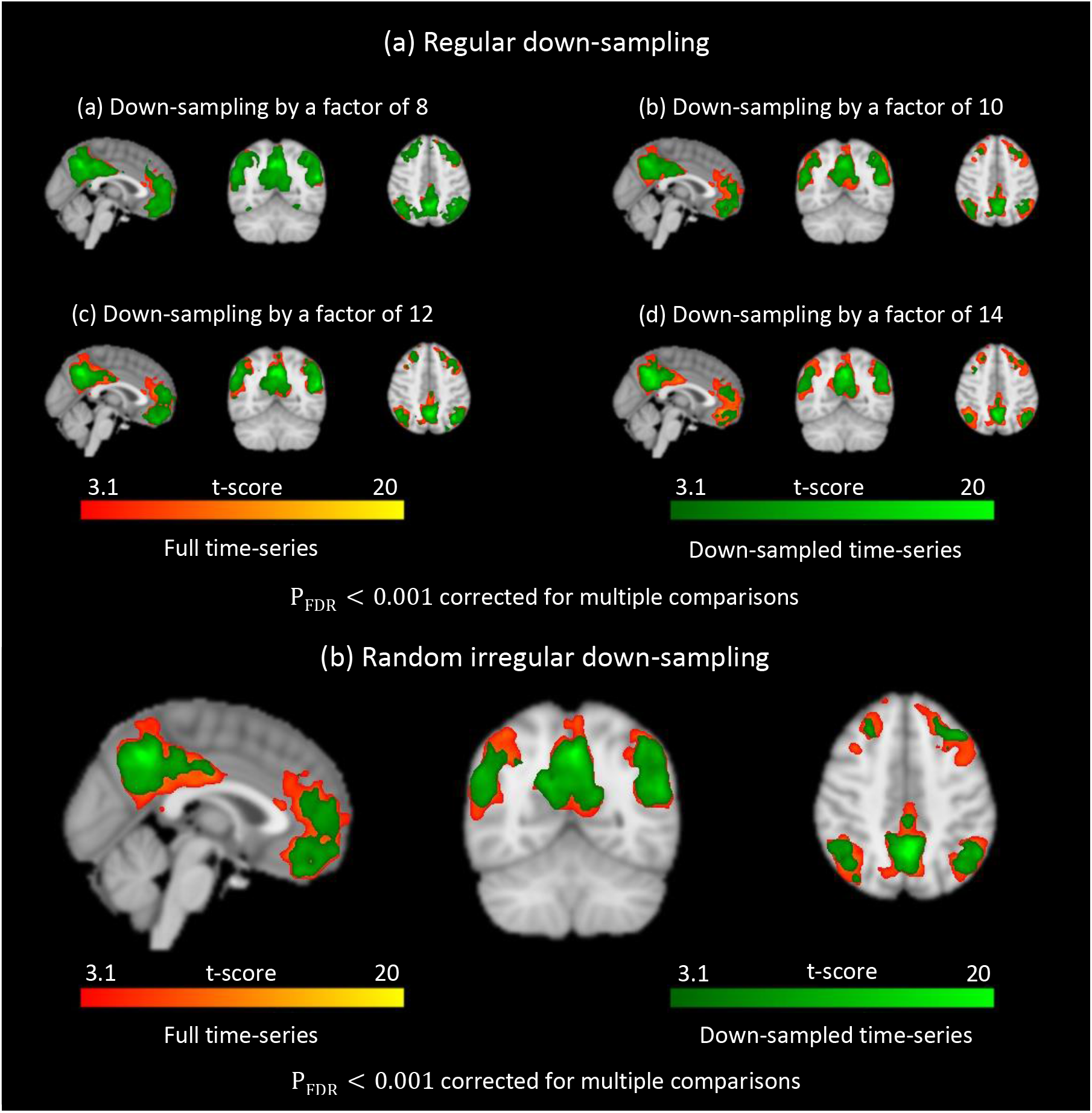
(a) Group-level analysis of seed-based correlations between individual voxels and a seed selected in the precuneus cortex (PCC). Correlations were calculated after regular down-sampling by various fixed down-sampling factors were performed. The values of the down-sampling factors were selected based on the 25th and 75th percentile of the boxplots shown in Figure 3. In all cases, the default mode network (DMN) was revealed. (b) Similar results as in (a) but this time down-sampling was performed using random irregular sampling intervals, which were obtained from a uniform distribution. The minimum and maximum values of the distribution were defined based on the 25th and 75th percentiles of the boxplots shown in Figure 3. In each case, the DMN was revealed from BOLD samples that did not necessarily coincide with large peaks in the BOLD time-series.

Figure 4b shows the same result, but in this case down-sampling was performed using random irregular sampling intervals between consecutive BOLD samples. Random sampling intervals were determined based on a uniform distribution. The minimum and maximum values of the distribution were defined based on the 25th and 75th percentiles of the boxplots shown in Figure 3. The result suggests that samples obtained from the BOLD signal, which may not coincide with large amplitude peaks, comprise relevant information that is sufficient to describe its slow dynamics observed during resting-state conditions.

Figure 5 shows conditional rate maps obtained using the residual time-series that resulted after regressing out the events corresponding to high amplitude peaks in the original BOLD time-series. They suggest that the spatial and temporal distribution of events defined at lower amplitude peaks in the BOLD is similar to the spatial and temporal distribution of events defined at higher amplitude peaks. This also confirms that important information to describe the slower dynamics of resting-state activity is not concentrated only in the higher amplitude peaks of the BOLD signal.

**Figure 5.**
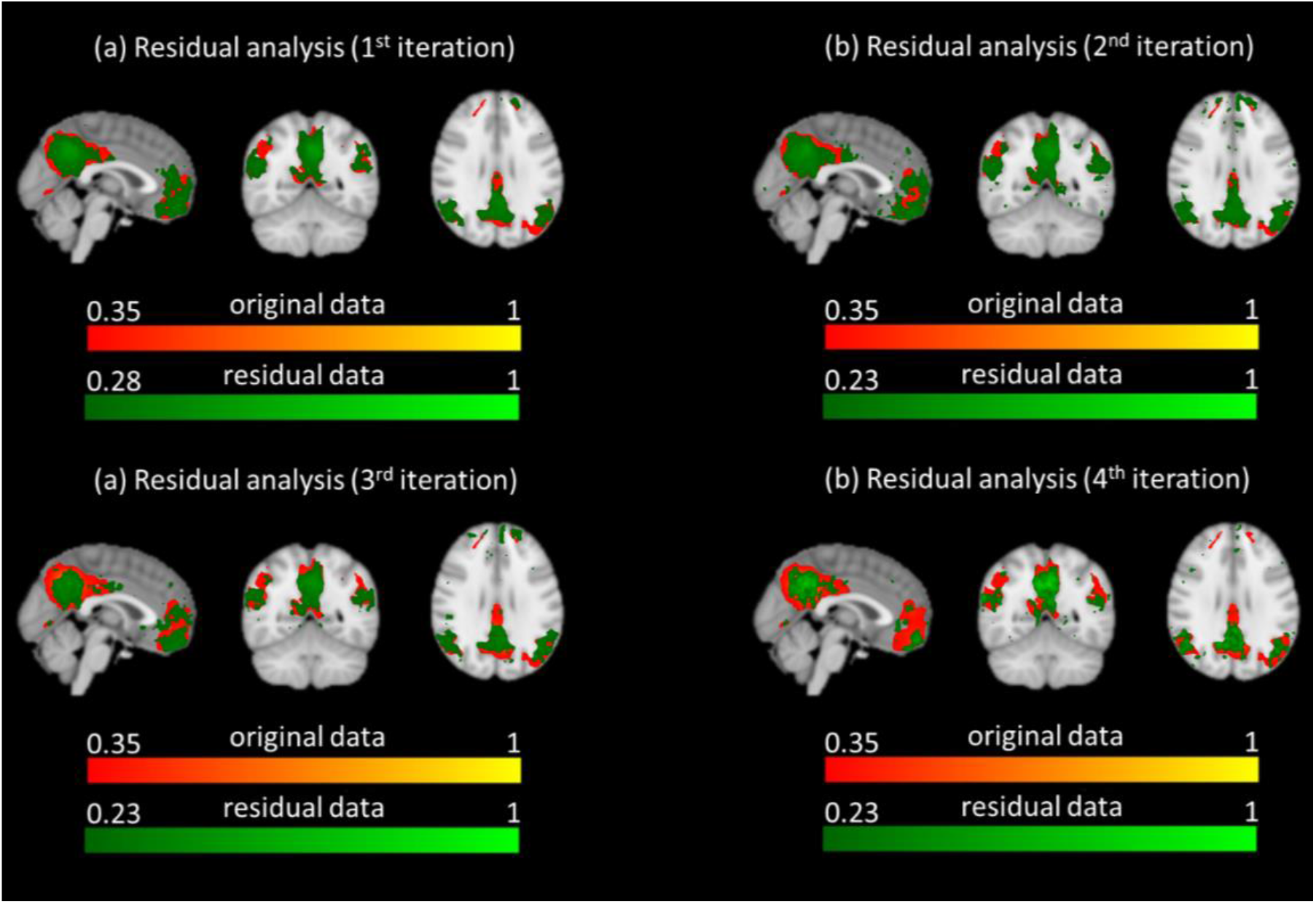
Group average conditional rate maps obtained across subjects using PPA events defined at the timing of lower amplitude peaks in the BOLD. The lower amplitude peaks were detected on the residual time-series after regressing out the higher amplitude peaks from the data. This procedure was repeated for four times: the first time PPA events were defined based on the original BOLD measurements. The other three times, PPA events were defined based on the residual time-series obtained in the previous iteration. In each case, the average conditional rate maps across subjects obtained from the residual time-series resembled the DMN. This suggested that even lower amplitude peaks of the BOLD signal convey important information regarding resting-state brain activity.

### 3.2. Convolutional sparse coding analysis

Figure 6 illustrates representative examples of CSC atoms obtained from the analysis of dataset 1. Figure 6a shows atoms that were classified as artifacts, which remained in the data after preprocessing. The assessment of the components as artifactual was based on evaluation of both the spatial patterns as well as signal morphology (temporal waveform). Starting from the left, the first atom was evaluated as a gradient artifact due to the high frequency peaks in its temporal pattern that possibly resulted from the fMRI gradient switchings. The second and third atoms correspond to ballisto-cardiogram and eye-blink artifact topographies that are typically observed in EEG-fMRI data.

**Figure 6.**
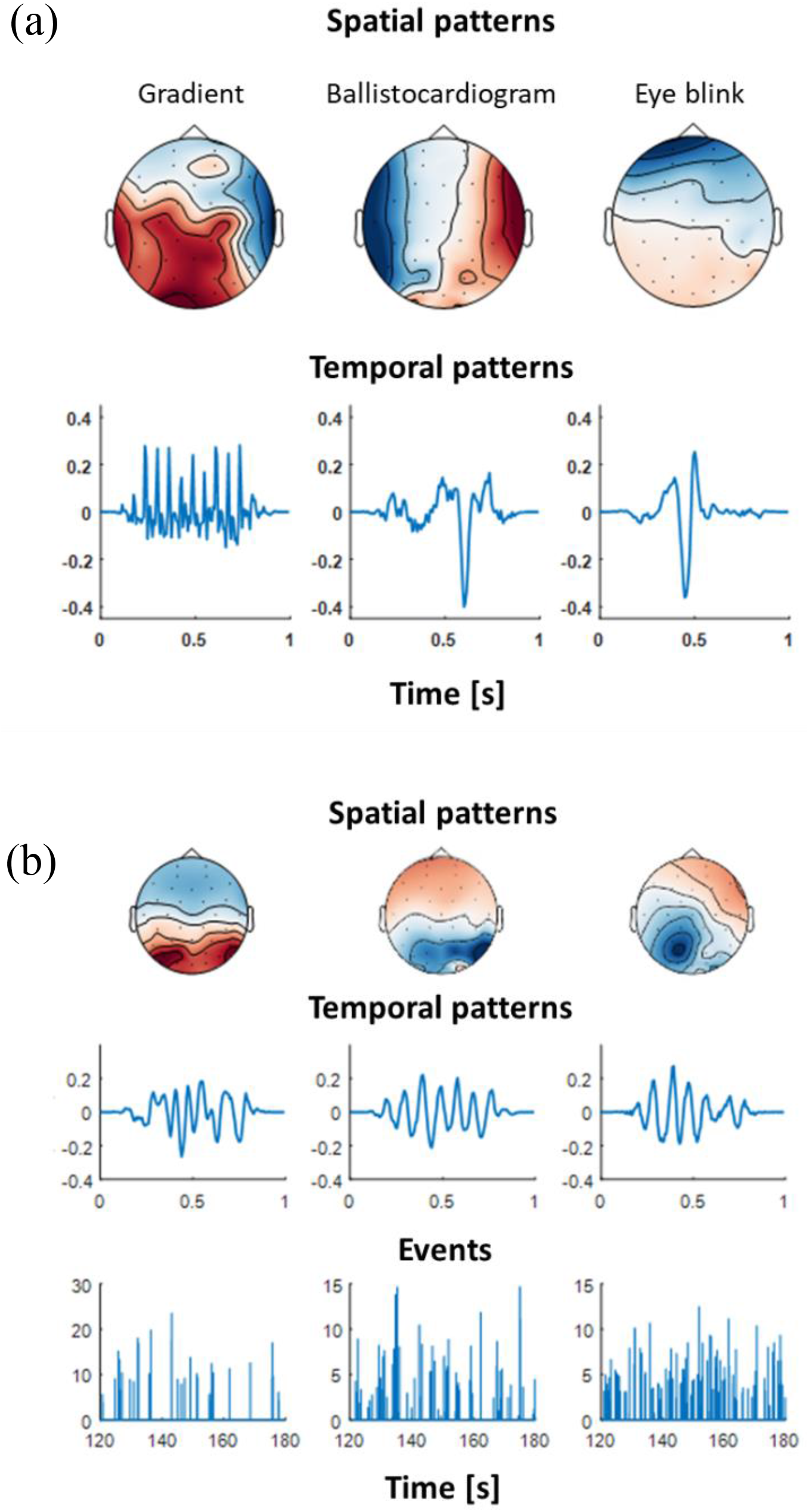
(a) Representative examples of atoms corresponding to artifacts in the EEG data. Assessment was based on both spatial patterns (top row) as well as signal morphology (bottom row). Starting from the left, the first atom was evaluated as gradient artifact due to the high frequency peaks in its temporal pattern that result from the fMRI gradient switchings. The second and third atoms correspond to typical ballisto-cardiogram and eye-blink EEG topographies. (b) Representative examples of atoms corresponding to brain activation during execution of the visual target detection task. Top row: CSC topographies showing fronto-parietal and parietal activation patterns. These patterns are consistent with activation of cortical attention and visual networks. Middle row: temporal pattern of each CSC atom. Bottom row: representative 20 s of sparse event timeseries associated with onset timing and magnitude of the temporal pattern of each atom.

Figure 6b shows atoms corresponding to brain activation associated with the visual target detection task (dataset 1). The CSC topographies show frontoparietal and parietal spatial patterns. These atoms are consistent with activation of the visual and cortical attention, which are engaged during the execution of the task. Atoms corresponding to brain activation associated with resting-state and motor task execution (dataset 2) are shown in Figure S1 and Figure S2 in the Appendix. Figure S1 shows the temporal waveform of a representative CSC atom as it appears in a segment of the original EEG sensor time-series. Figure S2 shows the spatial pattern, temporal waveform, as well as the power spectral density of the same prototypical CSC atoms shown in Figure S1. During resting-state, the temporal waveform shows higher power in the alpha (8-12) band, whereas during the motor task, the temporal waveform shows higher power in the and alpha and beta (>15 Hz) band.

### 3.3. Brain activation explained by CSC events

Figure 7a shows the results of the voxel-wise analysis performed using dataset 1. Activation maps obtained using CSC events detected in the EEG data are shown in green. Activation maps obtained with the subjects’ behavioral response time (RT) events, which were indicated by a button press, are shown in red/yellow. The results revealed activation in the lingual gyrus, insular cortex, left pre-central gyrus, and cingulate gyrus for both event types. In addition, CSC events revealed activation in the paracingulate gyrus. Figure 7b show group level activation maps obtained for the motor task data (dataset 2). In this case, red/yellow corresponds to the activation obtained using the subjects’ hand grip strength. The results revealed strong activation in the left pre-central gyrus (M1) for both event types. For both tasks, although the extend of activation obtained using the subjects’ behavioral response to each task is more widespread as compared to using the CSC events, overall there is concordance in the activation maps obtained in each case, suggesting that CSC events can be used to detect events associated with each task.

**Figure 7.**
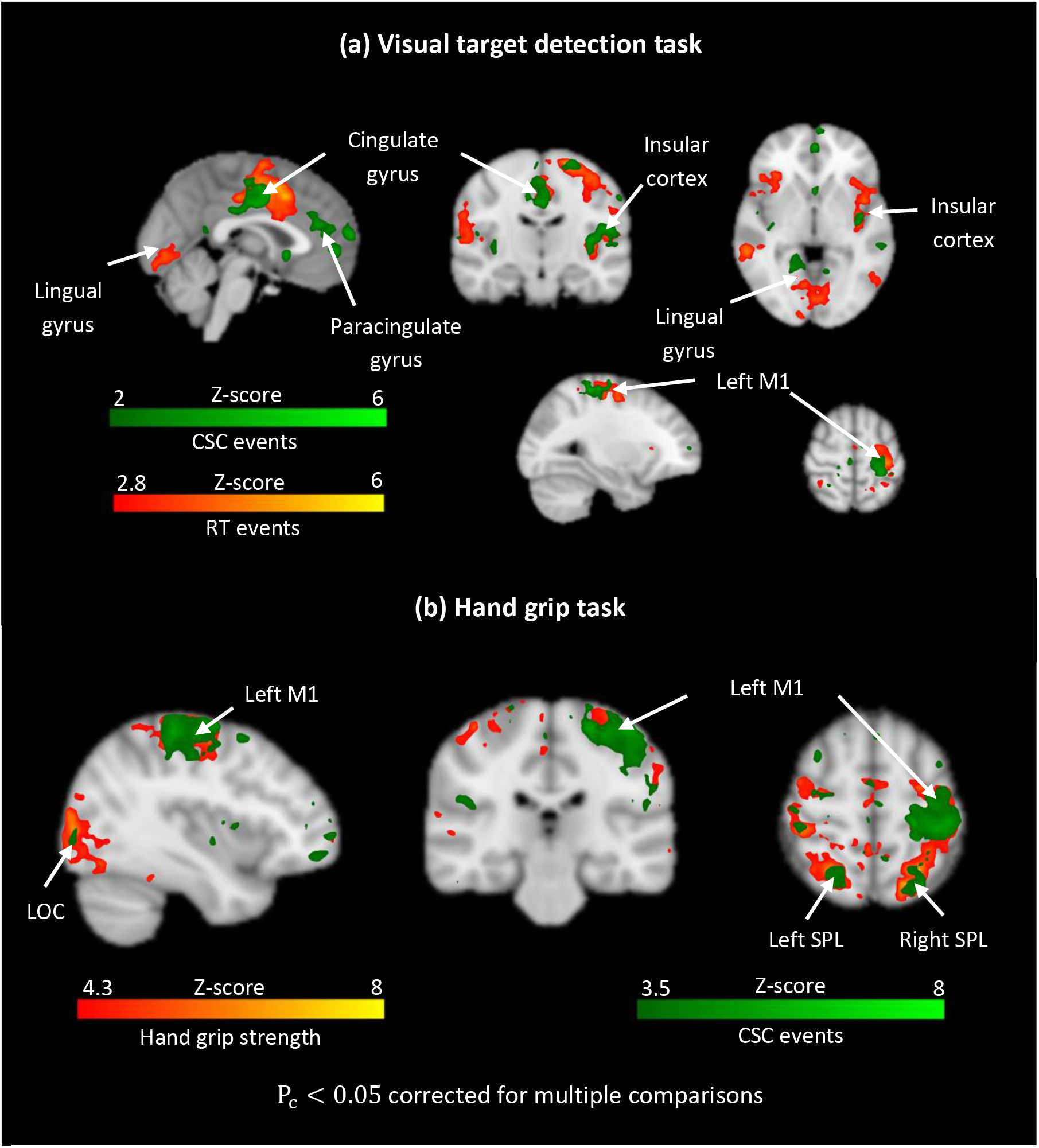
(a) Voxel-wise analysis showing brain activation at the group level during execution of a visual target detection task (dataset 1). Activated regions obtained from events detected in the EEG using CSC analysis are shown in green. Activated regions obtained using the subjects’ response time (RT) to the visual targets are shown in red/yellow. The maps show activation in the lingual gyrus, insular cortex, left pre-central gyrus (M1) and cingulate gyrus for both event types. CSC events also reveal activation in the paracingulate gyrus. (b) Group-level activation maps obtained during execution of a hand grip task (dataset 2). Activated regions obtained from events detected in the EEG using CSC analysis are shown in green. Activated regions obtained using the subjects’ hand grip strength are shown in red/yellow. The maps show strong activation in the left pre-central gyrus (M1) for both event types. In both conditions, although the subjects’ behavioral response to each task show more wide-spread activation, overall there is considerable overlap between the activation maps obtained with each method, suggesting that CSC is able to successfully detect neural events associated with each task.

Group average HRF estimates obtained during the motor task, as well as under resting conditions are shown in Figure 8. These estimates correspond to large functionally defined ROIs in which CSC events explained a large fraction of the variance in the BOLD signal. The ROIs included the left pre-central and superior parietal lobule cortices, for both experimental conditions. The HRF estimates obtained in each case exhibited consistent shapes, for most of the subjects. Representative BOLD signal predictions obtained for the left superior parietal lobule cortex are also shown in the same figure, for each experimental condition. They suggest that CSC events can be used to describe the slow oscillations in the BOLD under both conditions with some exceptions when BOLD exhibits large amplitude peaks that are not predicted by the EEG, which are possibly related to physiology or head motion.

**Figure 8.**
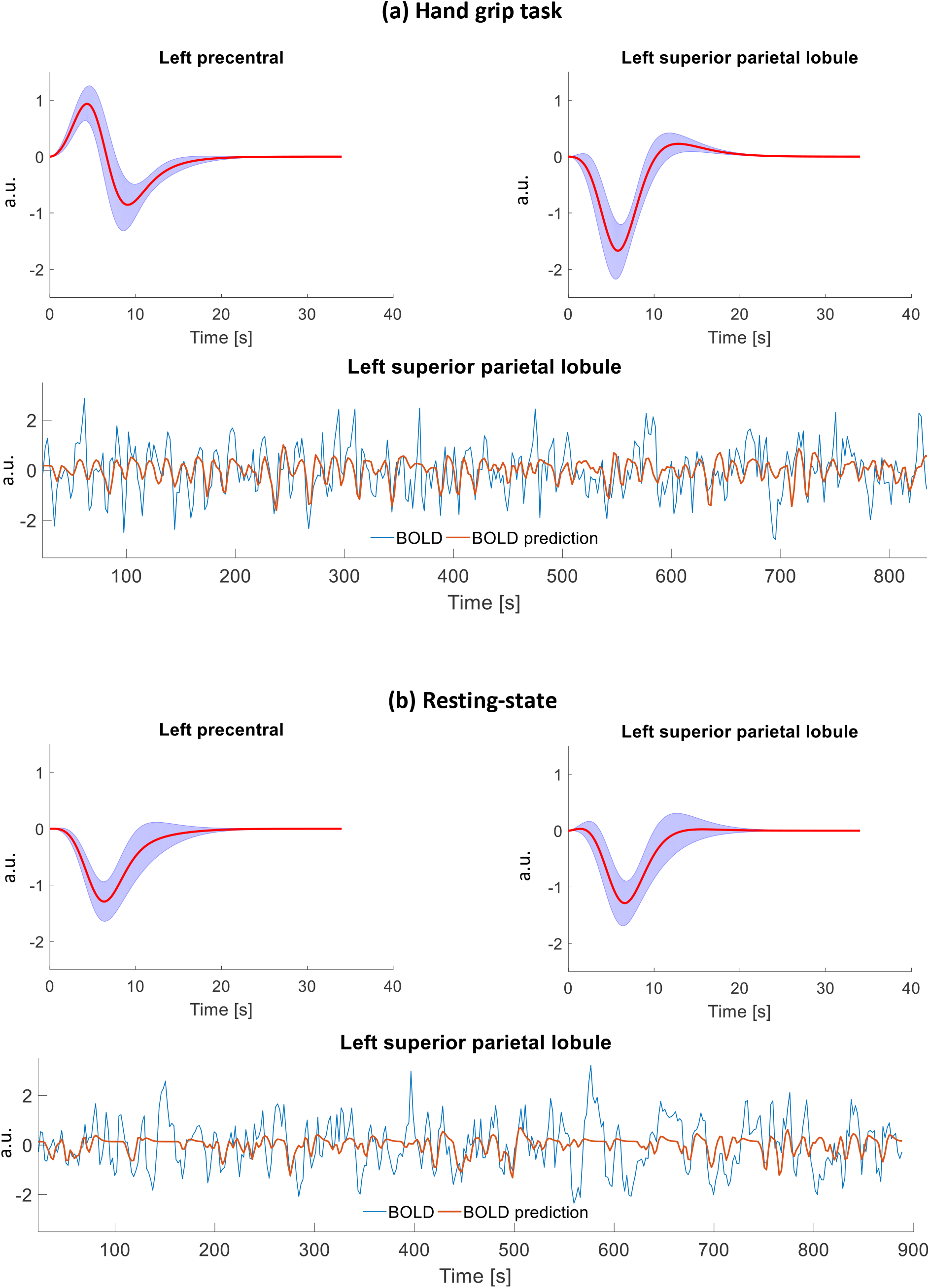
(a) Group average HRF curve shapes obtained in the left pre-central and left superior parietal lobule ROIs during motor task. The red curve corresponds to the mean HRF curve across all subjects. The blue shaded area corresponds to the standard error. The ROIs were functionally defined based on regions where EEG explained a large fraction of the variance in the BOLD signal (Figure 7b). Representative BOLD prediction in the left superior parietal lobule is shown in the lower panel. (b) Group average HRF curves obtained in the same ROIs under resting conditions. The ROIs were functionally defined based on regions where EEG explained a large fraction of the variance in the BOLD signal (Figure 7b). Representative BOLD prediction in the right occipital cortex obtained from the same subject as in (a) is shown in the lower panel.

Figure. 9 shows group level activation maps obtained using the resting-state data from dataset 2. Brain activation was evaluated using events defined in the EEG data with CSC along with event-related fMRI analysis. The results reveled widespread activation spanning multiple cortical areas.

**Figure 9.**
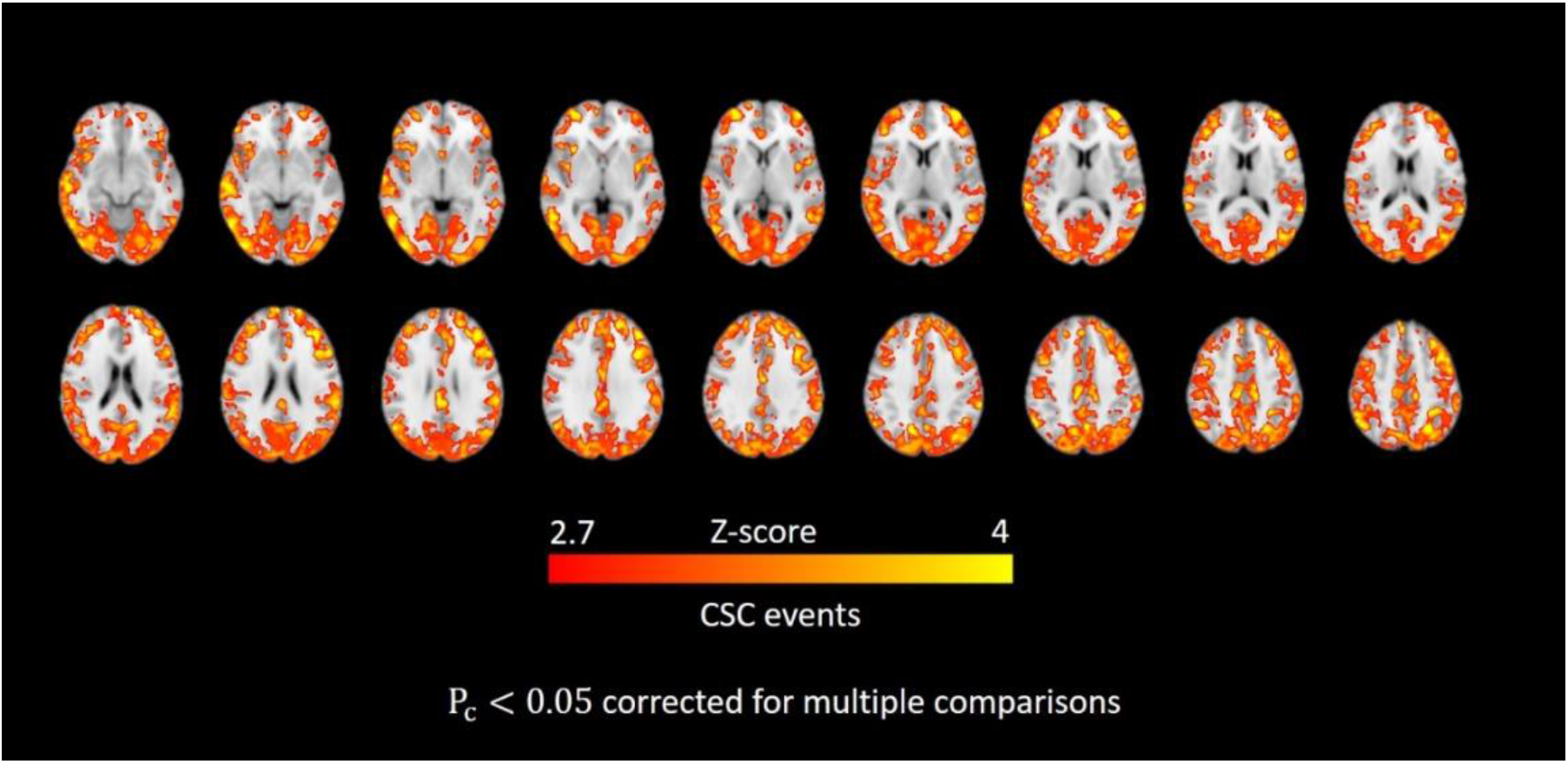
Voxel-wise analysis showing brain activation at the group level during resting-state. Brain activation was evaluated using CSC events and event-related fMRI analysis. The results reveled widespread and spanning multiple regions across the cerebral cortex.

## 4. Discussion

### 4.1. General

In this study, we investigated whether sparse, transient events detected in EEG data collected simultaneously with BOLD-fMRI can be used to describe the slow oscillations in the BOLD signal and to obtain reliable estimates of the HRF during task execution (visual target detection and a hand-grip task), as well as under resting-state conditions. This investigation was performed in light of the unfolding debate as to whether neural activity consists more of transient bursts of isolated events rather than rhythmically sustained oscillations (van Ede et al., 2018). To define events in the EEG we employed CSC analysis with rank-1 constraints (Jas et al., 2017; La Tour et al., 2018). We initially performed this analysis using the data collected during the two tasks, where events detected in the EEG were expected to explain higher BOLD signal variance in brain regions associated with each task. To model the BOLD signal, we employed FIR system analysis. Our results revealed activation in the lingual gyrus, insular cortex, left pre-central gyrus, and cingulate gyrus for the visual target detection task, and extensive activation in the left pre-central gyrus for the hand-grip task in accordance with previous studies in the literature (Sclocco et al., 2014; Walz et al., 2013; Xifra-Porxas et al., 2019). The same regions were also found to be activated using external measurements of the subjects’ behavioral response to each task. This suggested that CSC analysis can be used to detect reliable events in task-based EEG, which are associated with the task. It also suggested that sparse, transient events comprise relevant information that can be used to describe the slow fluctuations observed in the BOLD signal.

Subsequently, we performed the same analysis using resting-state data. Our results revealed that events detected in the EEG can be also used to explain the slow oscillations in the BOLD signal observed during the resting-state, despite the lower SNR associated with the later condition. They also suggested that CSC events can be used to obtain reliable HRF estimates, which exhibited consistent shapes across subjects. This line of research has important implications for the study of effective connectivity (Gao et al., 2016; Iwabuchi et al., 2014; Palaniyappan et al., 2018; G.-R. Wu et al., 2013; G. R. Wu et al., 2013) or functional connectivity (Gitelman et al., 2003; McLaren et al., 2012; Rangaprakash et al., 2018; Yan et al., 2018), as resting state HRF estimates are important in order to account for the hemodynamic blurring in BOLD-fMRI data.

### 4.2 Lower-amplitude co-fluctuations in BOLD-fMRI contribute to resting-state functional connectivity

Recent studies in the literature employed events defined at the timing of the large amplitude BOLD peaks to retrieve the HRF from resting-state fMRI data, which was subsequently used for hemodynamic deblurring (Abe et al., 2015; G.-R. Wu et al., 2013). A key hypothesis of this blind deconvolution approach was that relevant information of resting-state neural dynamics is encoded into the high amplitude peaks of the signal, which can be unveiled by PPA (Tagliazucchi et al., 2012). Along these lines, other studies proposed using sparse promoting deconvolution to define events, such as parameter free mapping (PFM) (Caballero Gaudes et al., 2013) or total activation (TA) analysis (Karahanoğlu et al., 2013). Moreover, similar studies proposed using related information associated to these events for the study of the spatial and temporal dynamics of resting-state brain activity. Such information included activation maps obtained using PFM event-related fMRI analysis (Petridou et al., 2013), clusters of fMRI frames obtained at the timing of PPA events (Liu et al., 2013; Liu and Duyn, 2013), or average frames obtained at the timing of high amplitude BOLD co-fluctuation (Betzel et al., 2019). While these works provided evidence that excluding these events results into a decrease in functional connectivity, they didn’t show that the structure of the resting-state functional networks also changes. The latter is an important index of coherent resting-state brain activity, which has been extensively used in the literature for quality control of resting-state fMRI preprocessing pipelines (Bright and Murphy, 2015).

In this work, using Pearson’s seed-based correlations with a seed selected in the PCC, we showed that voxels in the DMN co-fluctuate with the seed even when regular, or random, irregular down-sampling of the BOLD signal has been performed. In this case, samples obtained from the BOLD did not necessarily coincide with the high amplitude peaks of the signal, and yet the co-fluctuations of these samples between different voxels were found to preserve the spatial specificity of the network (Figure 4). This suggested that the relevant information of resting-state brain dynamics may not be condensed only in the high amplitude BOLD peaks, and that important information is also distributed in lower amplitude peaks. To investigate this further, we initially regressed the high amplitude peaks out from the original BOLD time-series in order to bring the lower amplitude peaks of the signal to the foreground. Subsequently, we applied PPA on the residual data and constructed seed-based conditional rate maps with a seed selected in the PCC. The results revealed that the derived conditional rate maps resembled the DMN (Figure 5), which confirmed that important information for resting-state brain dynamics is also concentrated in lower amplitude BOLD peaks.

These findings have important implications for the study of functional/effective connectivity using BOLD-fMRI, as well as for the study of HRF variability using events defined in the BOLD signal. They suggest that lower amplitude peaks convey important information about resting-state brain dynamics that should not be disregarded. Moreover, recent studies in the fMRI literature investigated a large number of fMRI preprocessing pipelines and pointed out that no pre-processing pipeline offers a perfect noise free signal (Parkes et al., 2018). Also, other studies showed that the network structure elicited by non-neural sources of BOLD signal variability is conformable to the structure of previously reported resting-state networks (Bright and Murphy, 2015; Chen et al., 2019; Nalci et al., 2019). On account of these considerations, we believe that the contribution proportion of neural versus non-neural sources in the high amplitude peaks of the BOLD cannot be easily elucidated. Hence, HRF or activity-inducing signal estimates obtained using blind deconvolution of the BOLD signal could be, to some extent, biased towards physiology processes or motion, and physiological interpretation of these estimates in terms of the underlying neural dynamics should be performed with caution.

### 4.3. HRF estimation using simultaneous EEG-fMRI data

In this work, we employed simultaneous EEG-fMRI data and CSC analysis with rank-1 constraints (Jas et al., 2017; La Tour et al., 2018) to define sparse events in the EEG that can be used to describe the slow fluctuations in the BOLD signal, as well as to obtain estimates of the unknown HRF. We believe that using EEG data acquired simultaneously with BOLD-fMRI is a more reliable approach to define neural-related events and brain states that can be used to describe the BOLD signal as (i) it provides more direct information with regards to neuronal activity, and (ii) this information is provided with a higher temporal resolution.

CSC analysis with rank-1 constraints is a recently developed dictionary learning technique for multi-channel EEG or magnetoencephalography (MEG) data decomposition into spatiotemporal atoms^3^. As it is illustrated by Figure 6, Figure S1, and Figure S2, this decomposition provides information with regards to accurate timing of events detected in the EEG, which is particularly important in event-related fMRI analysis. It also provides information with regards to the spatial pattern of brain activity, which can be used to descriminate between sources of brain activity for sources of non-neural origins in EEG data. Lastly, it also provides valuable information with regards to the morphology of the signals under consideration, which is important to better understant brain function under both health and desease (Cole and Voytek, 2019; Jones, 2016; Mazaheri and Jensen, 2008).

In general, CSC analysis assumes that neuronal activity detected with the EEG comes in packets or “bursts”, which only last for a few cycles. The waveform of these busts is considered to be time-invariant. This, however, is only an approxiation as the morphology of EEG signals is known to change over time (Amir and Gath, 1989; Li et al., 2011). This analysis generally offers better EEG signal approximations since the event waveforms are not constrained in narrow frequency bands (La Tour et al., 2018). In the present study, the selected CSC events revealed strong activations in parietal, occipital, and frontoparietal areas during both tasks. Moreover, most of the power of the oscillations of the waveforms associated with the selected atoms was found to be destributed in the alpha (8-12 Hz) and beta (15-30 Hz) bands, which is consistent with parietal and occipital activation due to visual stimulation employed in both tasks. In a recent work by (La Tour et al., 2018) CSC analysis was used for the analysis of MEG data collected during median nerve stimulation. The results revealed atoms characterized by waveforms oscilating in the mu band (~ 9 Hz) and localized activation in the primary somatosensory cortex. Overall, the results in the present study as well as in (La Tour et al., 2018) suggest that CSC analysis can be used to define events that are strongest in brain regions associated with the task.

Lastly, we note that even though CSC analysis revealed atoms with spatial patterns and temporal waveforms associated with known sources of noise (Figure 6a) it is likely that some noise (eg. BCG, head motion) has not been completely removed from the data during pre-processing and is still present in atoms deemed associated with neuronal activity. However, we believe that this has not affected our results (Figure 7Figure. 9), which revealed CSC atoms with high correlation with BOLD-fMRI in brain areas associated with the task.

### 4.4. Study limitations

The sparsity of the CSC event time-series 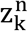 in equation (5) was controlled by the regularization parameter λ > 0. The value of this parameter was empirically defined for each dataset, aiming to balance between sparsity and BOLD prediction accuracy. We note that the selected value for this parameter is of the same order as the value selected for the analysis of the MNE somatosensory dataset (Gramfort et al., 2014, 2013) presented in (La Tour et al., 2018). Future work consists of extending the CSC algorithm employed herein to include an automated selection procedure for the parameter λ. This would make this technique more easily applicate to any dataset. Homotopy continuation procedures for sparse regression can be useful for that purpose (Efron et al., 2004; Osborne et al., 2000).

The present study was set out to investigate the underlying link between sparse neural events detected in the EEG with the contemporaneous changes in the BOLD signal. To this end, we employed EEG data, which were collected simultaneously with BOLD fMRI. Although this technique combines the excellent temporal resolution of the EEG with hemodynamic changes detected in high spatial resolution with BOLD-fMRI, it generally suffers from technical limitations associated with the EEG data acquisition inside the high magnetic field environment. Future work performed using optical imaging techniques, such as simultaneous EEG-FNIRS would help overcome these limitations.

## Conclusion

In this study, we initially employed seed-based correlations to show that samples obtained from the BOLD signal using various regular, as well as random sub-Nyquist sampling intervals, which did not necessarily coincide with large amplitude BOLD peaks, yield patterns of large scale resting-state neural dynamics observed with fMRI, such as the DMN. Subsequently, we performed a similar analysis using conditional rate mapping analysis. The results revealed that, in addition to the larger amplitude BOLD peaks, the spatial and temporal distribution of events defined at smaller amplitude BOLD peaks also resembles patterns of resting-state neural dynamics observed with fMRI. This suggested that using only events defined at the timing of the large BOLD amplitude peaks for HRF estimation, may yield biased estimates, which should be interpreted with caution.

To define more reliable neural events, we employed simultaneous EEG-fMRI data, along with CSC analysis. Our results suggested that the detected CSC events yield reliable activation maps obtained using event-related fMRI analysis. Our results also suggested that the events detected in the EEG yield consistent HRF estimates across subjects, even during resting-state conditions, where SNR is lower.

## Acknowledgements

This work was supported by funds from the Natural Sciences and Engineering Research Council of Canada (NSERC) Discovery Grants RGPIN-2019-06638 [GDM], and RGPIN-2017-05270 [MHB], the Fonds de la Recherche du Quebec - Nature et Technologies (FRQNT) Team Grant 254680-2018 [GDM], the Canadian Foundation for Innovation grant numbers 34362 [GDM] and 34277 [MHB]. The funders had no role in study design, data collection and analysis, decision to publish, or preparation of the manuscript.

## Appendix

**Figure S1.**
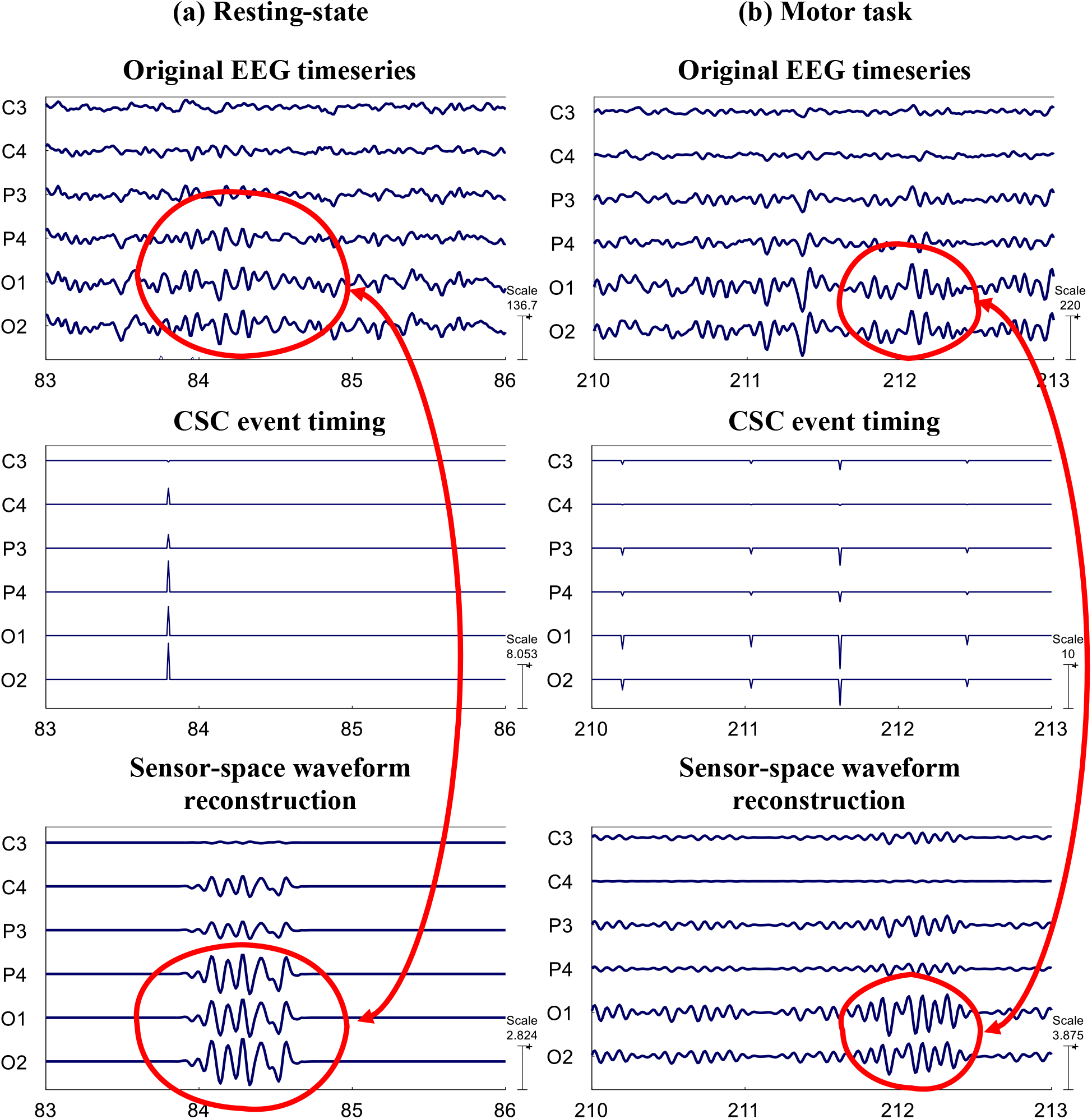
Representative CSC events corresponding to one atom detected in the EEG data of one subject during resting-state (left column) and motor task execution (right column). The topography and waveform of the detected atom are shown in Figure S2. Top row: original EEG time-series of 5 representative sensors. Middle row: CSC events. Bottom row: Waveform of the detected atom reconstructed at the sensor level.

**Figure S1.**
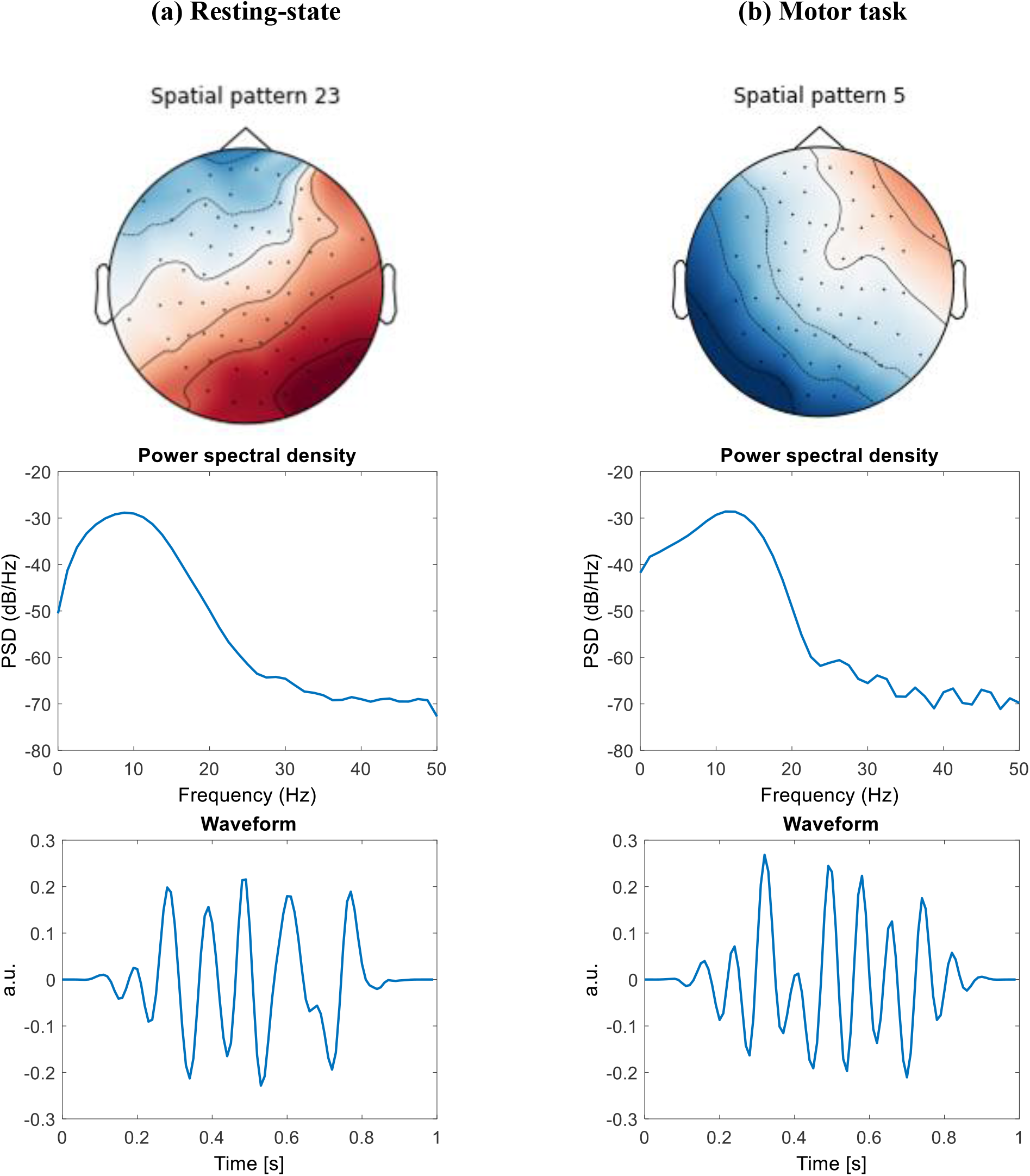
Representative CSC atoms detected in the EEG data of one subject during resting-state (left column) and motor task execution (right column) corresponding to the CSC events shown in Figure S1. Top row: Spatial patterns of brain activity associated with fronto-parietal activation. Middle low: Power spectrum density of the atom waveform showing high power in the alpha (8-12) band during resting state, and alpha and beta (>15 Hz) band during motor task execution. Bottom row: Prototypical CSC waveforms associated with the detected events shown in Figure S1.

## Footnotes

1 Dataset 1 was initially presented in (Walz et al., 2013). The dataset is publicly available through the OpenNeuro (Gorgolewski et al., 2017) online platform for sharing and analysis of neuroimaging data https://openneuro.org/datasets/ds000116.

5 The Harvard-Oxford cortical atlas is included in the FSL library (https://fsl.fmrib.ox.ac.uk/fsl/fslwiki/Atlases).

3 Open source code for convolutional sparse coding analysis of multivariate EEG/MEG data can be found at https://alphacsc.github.io/models.html.

## Notes

### Competing Interest Statement

The authors have declared no competing interest.

